# Multidimensional divergence supports parapatric speciation along an elevational gradient in ground-dwelling frogs on an oceanic island

**DOI:** 10.64898/2026.10.06.757056

**Authors:** Kier Mitchel E. Pitogo, Faben Hana Tesfazion, Aljohn Jay L. Saavedra, Ana Motta, Rafe M. Brown

## Abstract

Geographic isolation is a major driver of lineage divergence. However, strong ecological selection can promote reproductive isolation despite opportunities for gene flow in geographically continuous landscapes such as mountains. Here, we test whether ecological divergence across elevational gradients can promote speciation within a single mountain range. Integrating phylogenomic (ddRAD), morphological, and bioacoustic data, we test predictions of elevation-mediated parapatric speciation in *Platymantis* (*Lahatnanguri*) ground-dwelling frogs on Mindanao Island, Philippines. Phylogenomic analyses recovered an endemic radiation across neighboring mountain ranges and revealed a hierarchical pattern of diversification among three monophyletic lineages, with deeper divergence separating the mountain lineages Highland 1 and Highland 2 and more recent divergence occurring within a single mountain range. This topology identified a morphologically and acoustically distinct low-elevation lineage (<1,400 m a.s.l.) as closely related to the high-elevation Highland 1 lineage (>1,400 m a.s.l.), providing a natural test of divergence along an elevational gradient. The parapatric Lowland and Highland 1 lineages appear to have diverged relatively recently, as reflected in their shallow genome-wide differentiation, yet exhibit pronounced morphological and acoustic divergence. Population genomic analyses further recover discrete genomic clusters and no evidence of contemporary hybridization between the parapatric lineages, indicating independent evolutionary trajectories despite geographic continuity. Together, these findings are consistent with parapatric speciation along an elevational gradient and demonstrate how substantial phenotypic and reproductive-isolation divergence can accumulate despite limited genome-wide differentiation. More broadly, our results indicate that the exceptional biodiversity of oceanic archipelagos may arise not only from diversification among islands—and among mountains within islands—but also from lineage divergence along elevational gradients within mountains themselves. These empirical results support the biological plausibility of parapatric speciation within single mountain systems and highlight an underrecognized mechanism contributing to biodiversity in topographically complex oceanic archipelagos.

## INTRODUCTION

The exceptional biodiversity of oceanic islands is closely tied to questions about how lineages diversify and accumulate in geographically isolated and environmentally heterogeneous systems (Steinbauer et al. 2012; Brown et al. 2013; Triantis et al. 2015; Warren et al. 2015; Ali & Meiri 2019). Classical biogeographic models largely attribute this diversity to repeated colonization from neighboring landmasses followed by allopatric divergence in geographic isolation (MacArthur & Wilson 1963, 1967; Simberloff 1974; Whittaker et al. 2008). Subsequent work, however, recognized that speciation may also occur within islands, particularly on large, topographically complex landmasses where environmental heterogeneity can promote lineage divergence (Ogden & Thorpe 2002; Kisel & Barraclough 2010; Losos & Ricklefs 2009; Presgraves & Glor 2010). In such systems, elevational gradients, climatic transitions, and habitat discontinuities may generate opportunities for diversification even across geographically continuous landscapes.

Mountain systems are especially compelling for evaluating the possibility of *in situ* speciation because elevational gradients compress major environmental transitions across remarkably short geographic distances (Rahbek et al. 2019a,b). These steep ecological transitions can expose populations to contrasting selective pressures while maintaining opportunities for dispersal and gene flow, potentially driving population divergence (Keller et al. 2013; Guarnizo & Cannatella 2014; Funk et al. 2016; Rodríguez-Cajarville et al. 2022). When divergent selection across elevations is sufficiently strong to overcome the homogenizing effects of gene flow, reproductive isolation may evolve, providing a pathway to parapatric speciation (Bush 1975; Endler 1973; Templeton 1981; Rice & Hostert 1993; Gavrilets et al. 2000; Gavrilets 2014). Within a single mountain system, where populations are especially characterized by smaller range size, low local densities, and limited dispersal (Janzen 1967; Gaston 1996; McCain 2009), the chances for parapatric speciation is theoretically higher (Gavrilets et al. 2000; Gavrilets 2014). Although this mechanism is well supported by theoretical work in population genetics and speciation, compelling empirical examples remain relatively rare, particularly those integrating genomic, phenotypic, and behavioral evidence within a natural system (Caro et al. 2013; Linck et al. 2020).

Extensive field surveys in the Busa Mountain Range (BMR) of southern Mindanao Island, Philippines, documented morphologically and acoustically distinct ground-dwelling frogs in the Philippine-endemic genus *Platymantis* subgenus *Lahatnanguri* distributed across adjacent elevational zones (Pitogo et al. 2021a,b). This pattern of elevational structuring across a geographically continuous landscape raises two alternative explanations. First, lineage divergence may have occurred *in situ* along the elevational gradient itself, a plausible scenario given the extensive diversification documented in *Platymantis* (Brown et al. 2015; Deepak et al. 2025). Alternatively, these populations may represent secondary contact between lineages that diverged during periods of historical geographic isolation and later re-established contact within the mountain range. Distinguishing between these geographic pathways is challenging because alternative processes can produce similar phylogenetic (Via 2001; Losos & Glor 2003), genomic (Templeton 1981; Seehausen et al. 2017), spatial (Feder et al. 2013), and morphological patterns (Weber et al. 2021). However, integrating these independent lines of evidence offers a framework for evaluating predictions consistent with elevation-mediated parapatric speciation within a single mountain range (Lavrenchenko 2011; Nosil 2012; Caro et al. 2013).

Against this backdrop, the BMR provides a natural setting for testing whether ecological divergence along elevation can promote speciation within a geographically continuous mountain landscape. A critical first step in evaluating parapatric divergence is establishing whether the putatively divergent elevational lineages share a recent common ancestor. As expected under phylogenetic frameworks for within-region diversification (Moritz et al. 2000; Emerson 2002; Losos & Glor 2003), we first tested whether low-elevation (“Lowland”) and high-elevation (“Highland”) populations form a sister-lineage pair relative to other ground-dwelling *Platymantis* (*Lahatnanguri*) populations across the Mindanao biogeographic region to distinguish between secondary contact and *in situ* divergence within the BMR. Under a scenario of parapatric speciation, we predicted that these populations would be genomically less differentiated (Feder et al. 2013; Martin et al. 2013; Seehausen et al. 2017) yet exhibit independent evolutionary trajectories despite geographic proximity and opportunities for gene flow (Cruickshank & Hahn 2014)—a pattern also reflected in distinct genomic structures and demographic signatures (e.g., Feulner et al. 2015; Weber et al. 2021). If divergence was mediated by adaptation to contrasting elevational environments, we also expected corresponding shifts in morphology and acoustic mate-recognition signals (Smith et al. 1997; Orr & Smith 1998; Schluter 2001; Nosil 2012; Funk et al. 2016). Ultimately, we interpret concordance across genomic, morphological, and behavioral traits as evidence of an advanced stage of speciation despite recent divergence.

## MATERIALS & METHODS

### Fieldwork and sampling approach

Fieldwork was conducted by KMEP and AJLS from June to August 2024 within the Mount Busa Key Biodiversity Area, encompassing the Busa Mountain Range (BMR; Highland 1) and the Melibengoy Mountain Complex (MMC; Highland 2) (Figure 1A). Sampling in the BMR focused on both the northern slope (Highland 1a; 500–1800 m. a.s.l.) and the southern slope (Highland 1b; 200–2064 m. a.s.l.) to capture elevational and microhabitat variation and ensure representative sampling of *Platymantis* ground-dwelling frogs within the mountain range (Pitogo et al. 2020a,b). On the northern slope of BMR, we collected specimens of the Highland 1 lineage (KU 353024) and Lowland lineage (KU 353058) syntopically occurring at 1400 m. a.s.l. (identified based on call differences). Additional fieldwork was carried out in the MMC, where efforts targeted two peaks at approximately 1400–1600 m. a.s.l.. Individuals of Lowland and Highland lineages are initially identified based on body size and call differences, as initially reported (Pitogo et al. 2020a). Collected specimens were initially fixed in 10% buffered formalin, transferred to 70% ethanol for long-term preservation, and subsequently deposited in the University of Kansas Natural History Museum (KU) along with corresponding liver tissue samples preserved in absolute ethanol. Four specimens (KU 327594, KU 327609, KU 327596, KU 327598) from the geographically proximate Tampakan Highlands (Figure 1a), representing the widespread lowland lineage within southern Mindanao, were also included.

**Figure 1.**
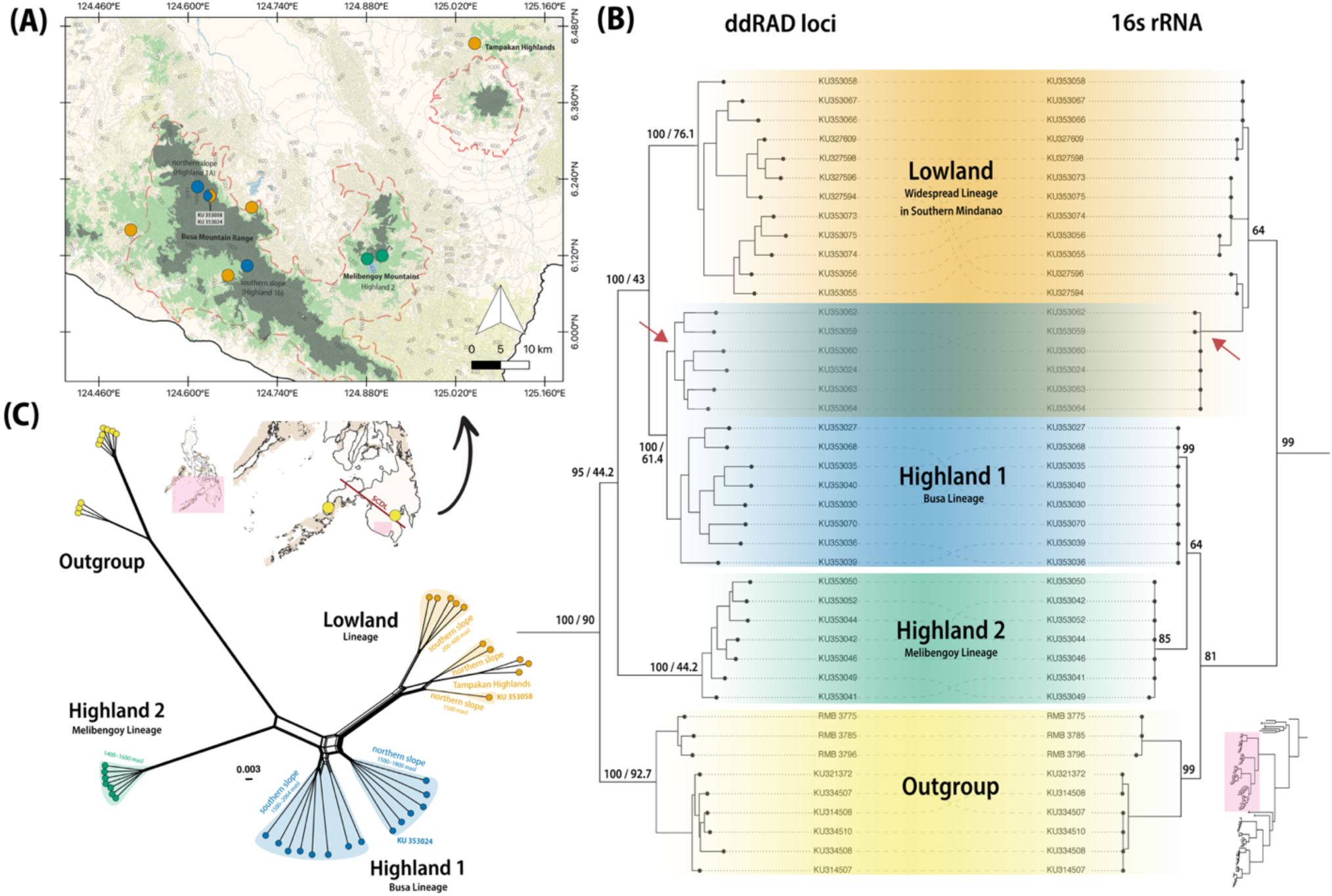
**(A)** Geographic distribution of focal lineages (Highland 1, Highland 2, Lowland, and Outgroup) in southern Mindanao, Philippines. The inset depicts the location of the study area within Mindanao Island. The South Cotabato–Daguma Lineament (SCDL), shown as a solid red line, delineates the hypothesized continental and oceanic portions of the island (Michaux 2012). **(B)** Phylogenetic relationships among focal lineages inferred from ddRAD loci (nodal support: ultrafast bootstrap values [UFBS] / site concordance factors) and mitochondrial 16S rRNA (UFBS), showing patterns of mito-nuclear discordance (red arrows indicate specimens that cluster with different lineages across the two datasets). The inset phylogeny shows that the focal lineages form a monophyletic group sister to all other ground-dwelling *Platymantis* (*Lahatnanguri*) from Mindanao. (C) Unrooted phylogenetic network of focal lineages illustrating relative genomic relationships among individuals and lineages. Colors are consistent in all figures and represent lineages as follows: Highland 1 (blue), Highland 2 (green), Lowland (orange), and Outgroup (yellow).

### Molecular data and bioinformatic processing

To identify major clades, we first conducted mitochondrial 16S sequencing across all *Platymantis* (*Lahatnanaguri*) ground-dwelling frogs from the Mindanao biogeographic region to guide our selection of representative samples for subsequent double-digest restriction-site associated DNA (ddRAD) sequencing (i.e., specimens that form a clade with each of the focal lineages). Genomic DNA was extracted and purified by lysing tissues with 20 µL of Proteinase K, followed by Solid Phase Reversible Immobilization (SPRI) bead extraction protocol (Roland & Reich 2012). For the 16S gene, a polymerase chain reaction (PCR) was conducted on a C1000^TM^ (Bio-Rad Laboratories, Inc.) thermal cycler to amplify ∼800-1000 base pairs (bp) of the mitochondrial 16S gene using forward primer (16Sc: GTR GGC CTA AAA GCA GCC AC) and reverse primer (16Sd: CTC CGG TCT GAA CTC AGA TCA CGT AG). The PCR conditions followed existing protocols used for Philippine amphibians (Brown et al., 2015). Amplification success was verified via electrophoresis on a 1.5% agarose gel. The PCR products were then purified and submitted for cycle sequencing and nucleotide sequence determination through standard protocols at Functional Biosciences®.

Double-digest RAD (ddRAD) libraries were prepared following published protocol (Peterson et al. 2012). Genomic DNA extracts were standardized to a concentration of 20 ng/µL and digested using the restriction enzymes *SbfI* and *MspI* (New England Biolabs). Digested fragments were ligated with unique P1 inline barcodes and P2 adapters. Samples were then pooled into four sublibrary batches, each assigned a unique i7 index to allow for multiplexing. These pools were size-selected for a 487–595 bp target using a 2% agarose cartridge on a Blue Pippin (Sage Science, Beverly, MA). Following PCR amplification and bead purification, the sub-libraries were sequenced in a single lane of an Illumina NovaSeq X Plus (150-bp paired-end reads) at Novogene (San Diego, CA).

Raw data were demultiplexed into sublibraries by Novogene and resolved into individual samples using the process_radtags module in Stacks v2.41 (Catchen et al. 2011, 2013). To ensure data consistency and prevent downstream cstacks errors, we used fastp (Chen et al. 2018) to generate a uniform-length dataset, hard-trimming and filtering all reads to a fixed length of 140 bp. Quality control included a sliding window truncation (window size = 4; mean Phred ≥ 20), excluding reads with >10 N bases or >30% low-quality bases (Phred < 20). Final data quality was verified using FastQC (Andrews 2010). We optimized the number of recovered loci following the “r80” protocol (Paris et al. 2017), testing various combinations of the parameters - m (minimum stack depth), -M (mismatches within individuals), and -n (mismatches between individuals).

Following initial quality control, the R package SNPfiltR (DeRaad 2022) was used to filter genotypes, in line with best practices for reduced-representation datasets (O’Leary et al. 2018). Genotypes were filtered for a minimum depth of 3 and genotype quality (GQ) of 30. We retained only biallelic SNPs, removed individuals with >87% missing data, and applied allele balance filtering (0.25–0.75) and a maximum mean depth of 300 to reduce the influence of paralogous loci. Rare variants were retained using a minimum minor allele count of 1. We then explored a range of per-SNP completeness thresholds (75–95%) and evaluated their effects on sample clustering using PCA and t-SNE ordinations, as implemented in SNPfiltR (DeRaad 2022). These analyses indicated that clustering patterns were primarily driven by lineage identity and were robust to variation in completeness thresholds. Based on these initial explorations, we retained a 75% per-SNP completeness threshold, resulting in a final dataset of 49,110 SNPs with 42 samples and ∼12% missing data (hereafter referred to as “filtered dataset”). We further filtered variants to reduce linkage disequilibrium by applying a minor allele count filter of 2 and subsequently thinning the dataset to retain one SNP per 500 bp, thereby minimizing the inclusion of closely linked loci. This resulted in a final, putatively unlinked SNP dataset comprising 42 samples and 2,927 SNPs, with 13.69% overall missing data (hereafter referred to as the “thinned dataset”).

### Inferring phylogenetic relationships

We reconstructed the evolutionary relationships among the focal lineages (three ingroup lineages and a designated outgroup) using mitochondrial (16S rRNA) and nuclear (ddRAD) datasets; outgroup was chosen based on inferred closer relationship from a published phylogeny of Ceratobatrachidae, the family that includes *Platymantis* (Brown et al., 2015) and their shared distribution with the ingroup within the hypothesized paleo-island precursor of present-day Mindanao Island divided by the South Cotabato-Daguma Lineament (SCDL; Figure 1A) (Yumul et al. 2020; Michaux 2010). To ensure a robust framework, we included representative samples from all recognized species of *Platymantis* (*Lahatnanguri*; Brown et al., 2015) and other widespread ground frogs within the Mindanao biogeographic region in both datasets (Pitogo et al. 2026). This broad sampling confirms the monophyly of our focal lineages within the wider *Platymantis* (*Lahatnanguri*) radiation.

Mitochondrial 16S rRNA sequences were assembled and edited de novo using Geneious Prime® 2025.1.3 (Kearse et al. 2012) and subsequently aligned with the MUSCLE algorithm (Edgar, 2022). We inferred a Maximum Likelihood (ML) phylogeny in IQ-TREE v2.3.6 (Nguyen et al., 2015), with the optimal substitution model selected based on BIC, as determined by the built-in ModelFinder function (Chernomor et al., 2016). Statistical support for the resulting nodes was assessed via 10,000 ultrafast bootstrap (UFBS) replicates. Parallel to the mitochondrial analysis, we reconstructed a phylogenomic tree using the whitelisted nuclear loci (49,110 SNPs) exported from the Stacks populations module. This concatenated matrix was analyzed in IQ-TREE using the ModelFinder Plus (-m MFP) option to identify and implement the best-fit model of evolution. To evaluate potential discordance, we calculated site concordance factors (sCF) following Minh et al. (2020). For both datasets, UFBS values ≥ 95 were interpreted as strong evidence for monophyly (Minh et al. 2013). Furthermore, sCF values exceeding the 33.3% random-expectation threshold were considered indicative of a phylogenetic signal favoring the reference branch over alternative arrangements in a localized quartet topology (Minh et al. 2020).

### Inferring evolutionary independence

To evaluate evolutionary independence among focal lineages, especially the three ingroups, we quantified both relative and absolute genomic divergence. Under divergence with gene flow, measures sensitive to within-population variation (e.g., *F_ST_*) and those independent of it (e.g., *D_a_*) are expected to covary positively, reflecting coordinated genomic differentiation under restricted gene flow (Cruickshank & Hahn 2014). Using the filtered dataset including both variant and invariant sites, we calculated pairwise *F_ST_* and net nucleotide divergence (*D_a_* = *d_XY_* − [(π1 + π2)/2]) to account for ancestral polymorphism and estimate divergence accumulated since population separation. We then assessed the relationship between *F_ST_* and *D_a_* across pairwise comparisons among ground frogs in the *Platymantis* (*Lahatnanguri*) from the Mindanao biogeographic subregion using linear regression. Pearson’s *r* and *R²* were used to quantify the strength of this relationship and establish a clade-wide baseline against which divergence in focal lineages could be interpreted, with significance determined at α =0.05.

To move beyond clade-level averages and examine genomic signatures of divergence directly, we performed sliding-window analyses of weighted *F_ST_* (50 kb windows; 10 kb step size) in VCFtools v0.1.16 (Danecek et al. 2011). We focused on three comparisons: Lowland versus Highland 1 (i.e., BMR populations), Lowland versus Highland 2 (i.e., MMC populations), and Highland 1 versus Highland 2 as a highland baseline. Comparing these distributions allowed us to visualize shifts in genomic differentiation across geographic contexts and stages of the divergence continuum.

To determine whether genomic divergence is translated into discrete genomic clusters, we evaluated population structure using complementary multivariate and ancestry-based approaches. We performed Principal Component Analysis (PCA) on the thinned, unlinked dataset using glPca in adegenet (Jombart & Ahmed 2011) and estimated ancestry coefficients using sparse non-negative matrix factorization (sNMF) in LEA v3.3.2 (Frichot & François 2014). We evaluated K = 1–10 with 100 replicates per K and selected the optimal model using cross-entropy. Results from K = 3–5 were visualized to capture hierarchical partitioning of genomic ancestry and transitions from broader ancestral connectivity to contemporary lineage structure.

To further evaluate whether genomic clustering reflects independent evolutionary lineages rather than ongoing admixture, we examined patterns of allele sharing and contemporary hybridization among the three ingroup lineages. We quantified excess shared ancestry across the resolved topology using *D*-statistics (ABBA–BABA; Durand et al. 2011) and f-branch (*f_b_*) analyses implemented in Dsuite (Malinsky et al. 2021). Because recent divergence and geographic proximity may generate shared genomic signatures through either retained ancestral variation or contemporary gene flow, we additionally tested for hybridization between the parapatric populations within the BMR (Lowland and Highland 1 lineages). Using the R package triangulaR (Wiens et al. 2025), we identified ancestry-informative markers (Δ ≥ 0.90) and applied stringent thinning to minimize physical linkage (one SNP per scaffold; n = 91 variants). Together, these analyses tested whether the parapatric lineages maintain distinct genomic backgrounds despite geographic continuity and limited shared ancestry.

We also quantified demographic signatures across focal lineages to complement tests of genomic independence. Nucleotide diversity (π) was calculated at both individual and population levels, and individual heterozygosity was estimated in VCFtools v0.1.16 (Danecek et al. 2011) as the proportion of heterozygous sites across recovered loci. Sliding-window estimates of π and Tajima’s *D* were calculated using 50-kb windows with a 10-kb step size. Given the inferred transition into lowland habitats, we expected the lowland lineage to exhibit relatively elevated genetic diversity and more negative Tajima’s *D* values, patterns commonly associated with recent population expansion (Tajima 1989; Hewitt 2004). Demographic patterns were additionally visualized using an unrooted phylogenetic network in SplitsTree v4.19.1 (Huson & Bryant 2024).

Finally, we tested whether divergence associated with the transition of the Lowland lineage into lower elevations was accompanied by shifts in molecular evolutionary rates, as warmer environments have been hypothesized to increase rates of molecular evolution (Wright et al. 2010; Puurtinen et al. 2016). We performed Relative Rate Tests following the genomic framework of Pang et al. (2025). From an initial set of 3,015 ddRAD loci, we retained 2,887 loci represented across all four focal groups (Highland 1, Highland 2, Lowland, and Outgroup). For each locus, a single representative sequence from each lineage was used to construct four-taxon alignments, and pairwise genetic distances were estimated under a Jukes–Cantor (JC69) model using the R package ape (Paradis & Schliep 2019). Statistical significance was assessed using 1,000 bootstrap replicates, in which loci were resampled with replacement and 95% confidence intervals were calculated for pairwise rate comparisons, particularly among the three ingroup lineages.

### Inferring morphological and acoustic divergence

Consistent with our evaluation of genomic independence, we next tested whether lineage differentiation was accompanied by morphological and acoustic divergence, as shifts in these traits may reflect the evolution of reproductive isolation. To characterize morphological differentiation among focal lineages, we measured 66 fluid-preserved adult male specimens (Highland 1, n = 19; Highland 2, n = 10; Lowland, n = 26; Outgroup, n = 11) supplemented with measurements from additional ground-dwelling *Platymantis* (*Lahatnanguri*) populations across the Mindanao biogeographic region (Pitogo et al. 2026). Following Watters et al. (2016), we quantified 20 standard amphibian mensural traits including snout–vent length (SVL), limb dimensions, and head and digit measurements. To minimize observer bias, all measurements were collected by KMEP from high-resolution digital images (Canon EOS 850D; 100-mm macro lens) using ImageJ (Rueden et al. 2017). Raw measurements were size-standardized using a group-level allometric growth model (function allom, type = “population1”) implemented in GroupStruct (Chan & Grismer 2022).

To evaluate the relationship between phenotypic and genomic divergence, we compared pairwise morphological distances with absolute genetic divergence (*d_XY_*) across all measured *Platymantis* (*Lahatnanguri*) populations. Morphological differentiation between population centroids was quantified using Mahalanobis distance (*D_M_*), calculated from pooled within-population covariance matrices to account for trait correlations. Pearson’s *r* and *R²* were used to quantify the strength of this association, with significance determined at α = 0.05. We additionally performed Linear Discriminant Analysis (LDA) using MASS (Venables & Ripley 2002) to identify combinations of traits that maximize lineage separation.

To complement morphology, we evaluated acoustic divergence using representative advertisement calls from each focal lineage; recordings were deposited in the Macaulay Library (Cornell Lab of Ornithology): Highland 1 (Accession nos. ML663479721, ML663479723), Highland 2 (ML663480135, ML663480134), Lowland (ML663481074, ML663479722), and Outgroup (ML663480830, ML663480833, ML663480835). From each recording, we selected a high-quality segment containing continuous calling activity and a high signal-to-noise ratio for analysis. Acoustic analyses and spectrogram generation were conducted using the R package seewave (Sueur et al. 2008). Temporal parameters, including note repetition rate and note duration, were extracted using the automated timer function with a standardized amplitude threshold (25%) and smoothing window (500 samples). Note repetition rate was calculated as notes per second and note duration as the mean width of detected notes.

Spectral analyses were performed using representative non-overlapping notes (Highland 1, n = 8; Highland 2, n = 7; Lowland, n = 6; Outgroup, n = 13). Notes were extracted using lineage-specific windows (typically 20–40 ms) matched to note duration and filtered using adaptive band-pass ranges (1–4 kHz) to minimize environmental noise. Mean power spectra were generated for each note, and dominant frequency was defined as the spectral peak with maximum relative amplitude. All peak detections were manually verified against spectrograms and power spectra to confirm correspondence with the primary resonant frequency.

## RESULTS

### Evolutionary history and topological discordance among lineages

The inferred topologies from both the mitochondrial 16S rRNA and ddRAD genomic trees recovered a monophyletic group containing the three ingroup lineages (Highland 1, Highland 2, and Lowland) and the designated outgroup (Figure 1B). This group is strongly supported as sister to the rest of the Mindanao ground-dwelling *Platymantis* frogs in the subgenus *Lahatnanguri (*16S UFBS = 99; ddRAD UFBS = 100, sCF = 90%). However, the two datasets exhibit mito-nuclear discordance in their internal relationships. In the mitochondrial 16S tree, the Lowland lineage is recovered as sister to a clade comprising Highland 1, Highland 2, and the Outgroup with strong support (UFBS = 99; Figure 1B). Within this group, Highland 1 and Highland 2 form a sister-pair (UFBS = 64) that is collectively sister to the Outgroup (UFBS = 81), although both relationships have moderate support.

In contrast, the ddRAD genomic tree recovers the Outgroup as sister to the three ingroup lineages with strong support (UFBS = 100; sCF = 90%; Figure 1B). Highland 1 and the Lowland lineages are recovered as reciprocally monophyletic sister groups (UFBS = 100; sCF = 43%), which together form a clade sister to Highland 2 (UFBS > 95; sCF = 44.2%). Notably, Highland 1 individuals collected from the northern slope of the BMR cluster within the Lowland clade in the mitochondrial 16s rRNA ML tree despite their well-supported nuclear affinity with Highland 1. Our results appear to provide support for the potential interpretation of mitochondrial introgression or incomplete lineage sorting.

### Genomic divergence and population structure across the speciation continuum

Genome-wide divergence across the ground-dwelling *Platymantis* (*Lahatnanguri*) clade from the Mindanao biogeographic region follows a predictable trajectory characterized by a strong coupling between absolute and relative genomic differentiation (Figure 2A). Across populations within the region, net nucleotide divergence (*D_a_*) and relative differentiation (*F_ST_*) were strongly positively correlated (Pearson’s *r* = 0.72, *R²* = 0.52, *p* < 0.01). This relationship was even stronger within the focal lineages (Highland 1, Highland 2, Lowland, and Outgroup; *r* = 0.90, *R²* = 0.81, *p* < 0.01). Despite this shared trajectory, the parapatric lineages (Highland 1 and Lowland) consistently occupied the lower end of both *D_a_* and *F_ST_* values relative to the clade-wide pattern, placing them at the earliest stages of the genomic divergence continuum within Mindanao ground-dwelling *Platymantis* (*Lahatnanguri*).

**Figure 2.**
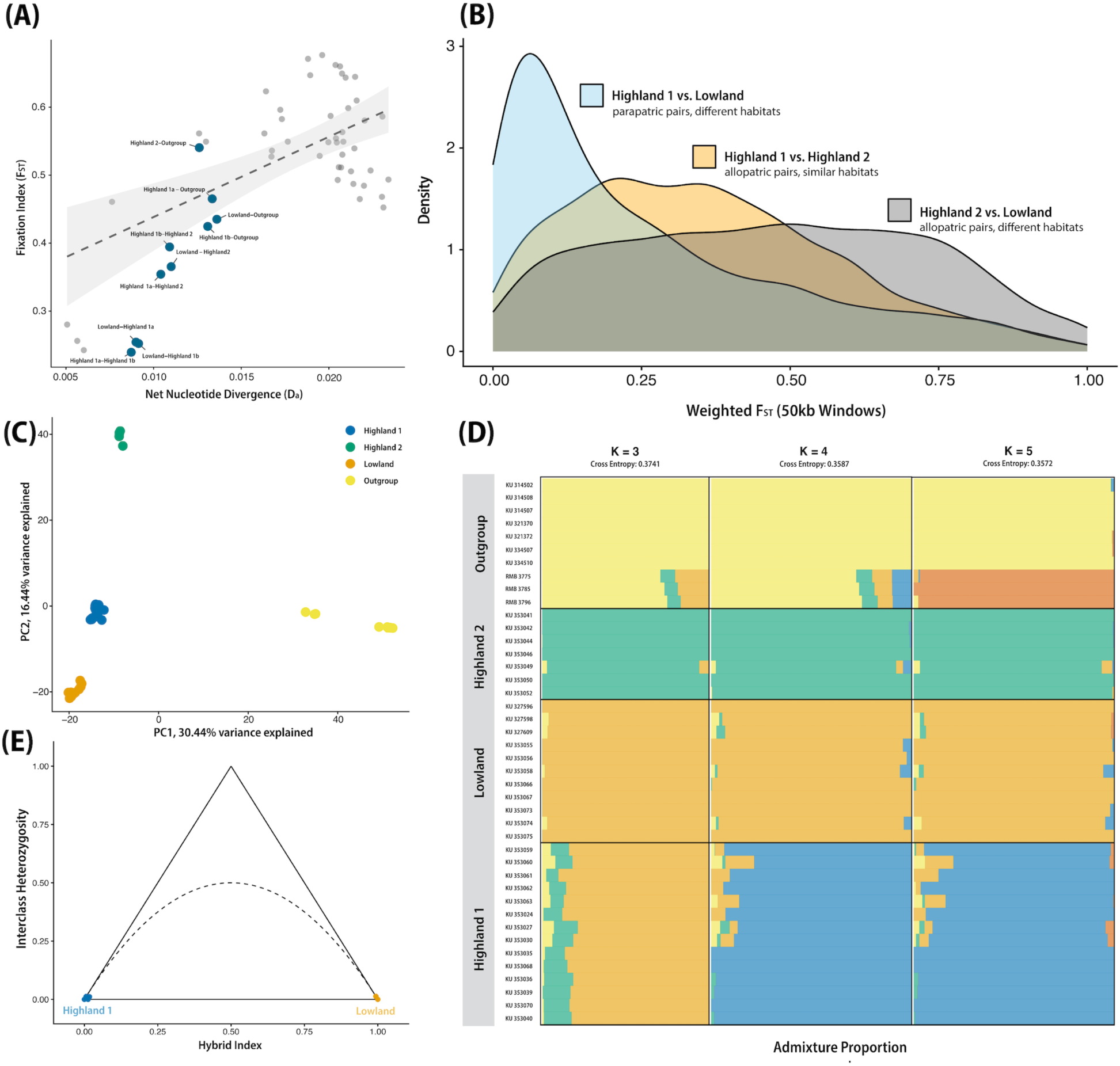
Genomic independence among focal lineages. **(A)** Genome-wide relative differentiation (*F_ST_*) plotted against net nucleotide divergence (*D_a_*), illustrating the positive relationship between these metrics. Blue points represent pairwise comparisons just among focal lineages, whereas gray points correspond to all ground-dwelling *Platymantis* (subgenus *Lahatnanguri*) populations across the Mindanao biogeographic region, including the focal group. **(B)** Density distributions of windowed *F_ST_* values, illustrating shifts in genomic differentiation across the speciation continuum and among geographic scenarios. **(C)** Principal Component Analysis (PCA) of genome-wide SNP data showing discrete clustering of focal lineages in multivariate space. **(D)** Ancestry coefficients inferred using sparse non-negative matrix factorization (sNMF) for K = 3–5, with K = 5 representing the optimal model based on the lowest cross-entropy score. **(E)** Triangle plot of interclass heterozygosity between the parapatrically distributed Highland 1 and Lowland lineages, showing no evidence of contemporary hybridization despite geographic continuity.

Patterns of genome-wide differentiation further support this interpretation. Density distributions of weighted *F_ST_* across genomic windows revealed clear shifts associated with divergence stage (Figure 2B). Comparisons between Highland 1 and Lowland produced a strongly left-skewed, leptokurtic distribution, indicating that most genomic regions remain weakly differentiated. In contrast, comparisons involving Highland 2 (Highland 2–Lowland and Highland 1–Highland 2) showed broader, more platykurtic distributions with greater variance across the genome. These patterns mirror the higher *D_a_* values observed among these comparisons (Figure 2A).

This genomic differentiation is reflected in discrete evolutionary groups. PCA recovered clear and non-overlapping genomic clusters corresponding to focal lineages (Figure 2C). The Outgroup separated along PC1 (30.4% variance explained), while PC2 (16.4%) further distinguished Highland 2 from Highland 1 and Lowland. Consistent with their shallow divergence, Highland 1 and Lowland occupied relatively proximate positions in multivariate space. Likewise, ancestry analyses recovered similar hierarchical structures. The sNMF analysis identified K = 5 as the best-supported model (cross-entropy = 0.3572; Figure 2D). At K = 3, Highland 1 and Lowland retained broadly similar ancestry components, whereas Highland 2 was already differentiated. By K = 4, all focal lineages formed distinct ancestry clusters. Notably, Highland 1 individuals sampled along the northern slope of the BMR exhibited minimal shared ancestry with Lowland despite continuous elevational distributions.

The recovery of discrete genomic groups despite shallow differentiation prompted further tests of whether this structure reflects ongoing gene flow or retained ancestral variation. D-statistics and f-branch analyses detected excess allele sharing between Highland 1 and Highland 2 (*D* = 0.355, *Z* = 5.51, *p* < 0.001; *f_b_* = 0.151), but not between Lowland and either highland lineage. Hybrid analyses further showed that parapatric populations maintained distinct genomic assignments with no evidence of intermediate ancestry (Figure 2E). Together, these results indicate that the excess shared ancestry detected between Highland 1 and Highland 2 is consistent with greater historical connectivity among high-elevation populations, whereas the Lowland and Highland 1 lineages maintain largely independent genomic backgrounds despite geographic continuity.

Demographic signatures varied markedly among focal lineages and further supported their genomic distinctiveness. Density distributions of nucleotide diversity (π) reveal substantial differences in standing genomic variation among lineages, with Highland 1 and the Lowland lineage exhibiting broader distributions shifted toward higher values relative to Highland 2 and the Outgroup (Figure 3A). This pattern is mirrored at the individual level, where Highland 1 exhibited the highest heterozygosity (mean = 0.064 ± 0.002), followed by the Lowland lineage (0.049 ± 0.007), Highland 2 (0.042 ± 0.002), and the Outgroup (0.033 ± 0.004) (Figure 3B). Sliding-window analyses of Tajima’s *D* further revealed contrasting demographic signatures between Highland 2 and the other lineages. Highland 1 showed the most negative mean Tajima’s *D* values (–0.209 ± 0.866), followed by the Lowland lineage (–0.068 ± 0.950) and the Outgroup (–0.081 ± 0.964), whereas Highland 2 exhibited predominantly positive values (0.591 ± 1.020) (Figure 3C).

**Figure 3.**
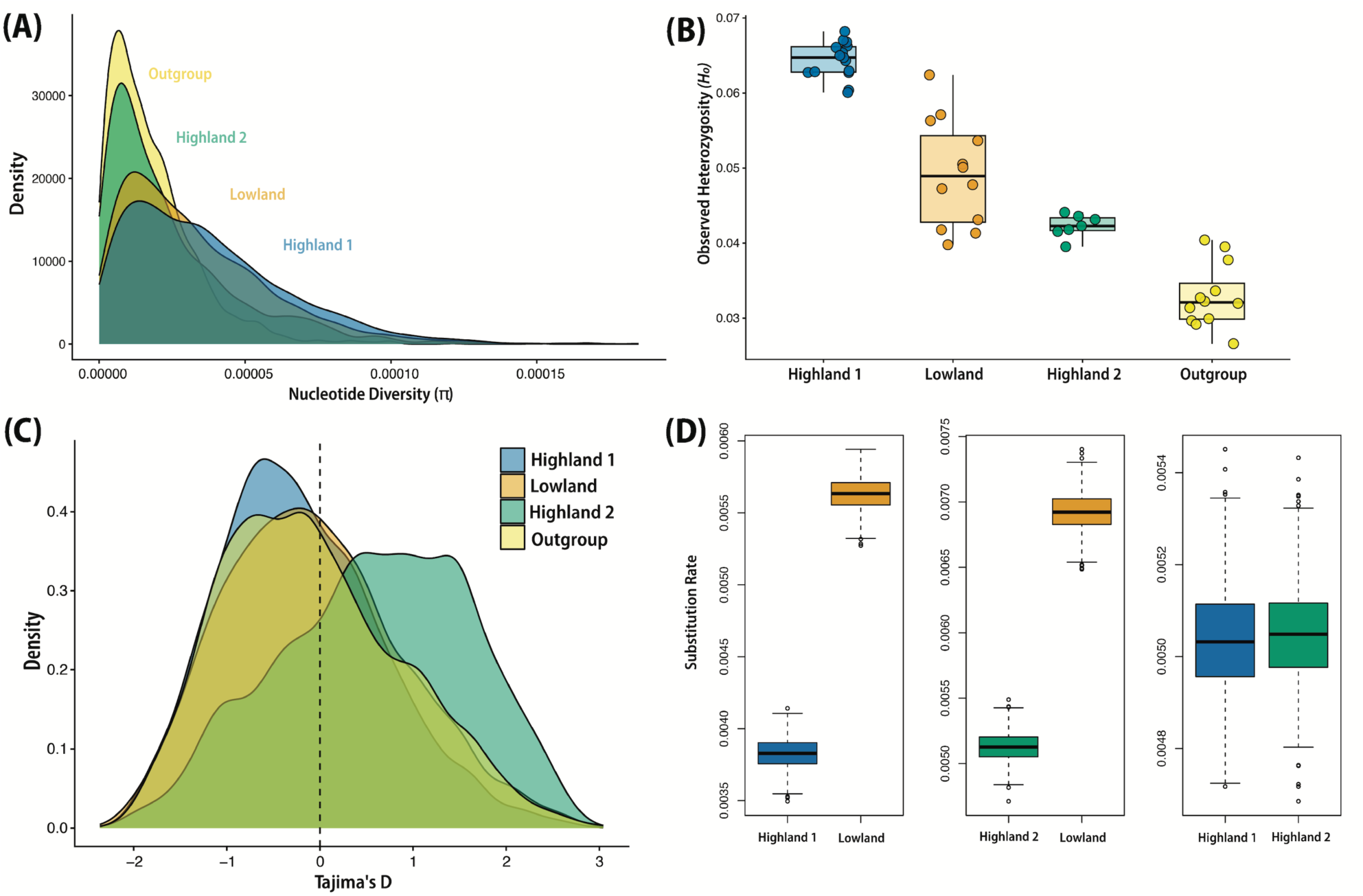
Demographic patterns inferred from genome-wide data among focal lineages. **(A)** Density distributions of population-level nucleotide diversity (π), showing elevated genomic diversity in Highland 1 and Lowland relative to Highland 2 and the Outgroup. **(B)** Individual-level heterozygosity estimates, reflecting similar patterns of genomic variation across lineages. **(C)** Density distributions of genome-wide Tajima’s *D*, demonstrating that Highland 2 exhibits a greater proportion of genomic windows with positive values, whereas the remaining lineages are characterized by predominantly negative values. **(D)** Pairwise substitution-rate estimates from Relative Rate Tests among ingroup lineages, showing elevated rates in the Lowland lineage relative to both highland lineages and minimal rate differences between Highland 1 and Highland 2.

Relative rate tests identified significant asymmetry in molecular evolutionary rates involving the Lowland lineage (Figure 3D). Pairwise comparisons recovered elevated substitution rates in Lowland relative to both Highland 1 (rate asymmetry = 0.320) and Highland 2 (rate asymmetry = 0.259), whereas the two highland lineages exhibited negligible rate differences (rate asymmetry = 0.002). Overall, the Lowland lineage evolves up to ∼38% faster than the highland lineages, representing a rate disparity approximately six-to seven-fold greater than that separating Highland 1 and Highland 2. Together, these results indicate that the Lowland lineage is characterized by elevated molecular evolutionary rates despite relatively shallow genome-wide differentiation from its sister Highland 1 lineage.

### Morphological and acoustic divergence

Morphological divergence does not scale predictably with genomic distance across recently diverged ground-dwelling populations of *Platymantis* (*Lahatnanguri*) from the Mindanao biogeographic region (Figure 4A). Relationships between Mahalanobis distance (*D_M_*) and absolute genetic divergence (*d_XY_*) were non-significant both across the broader clade (*r* = 0.10, *p* = 0.465) and within focal lineages (*r* = −0.02, *p* = 0.953), with genomic distance explaining little morphological variation (*R²* ≤ 0.01). Despite spanning only a narrow range of genomic divergence (*d_XY_* ≈ 0.011–0.018), focal lineages exhibited a wider range of morphological differentiation (*D_M_* ≈ 2.62–3.72).

**Figure 4.**
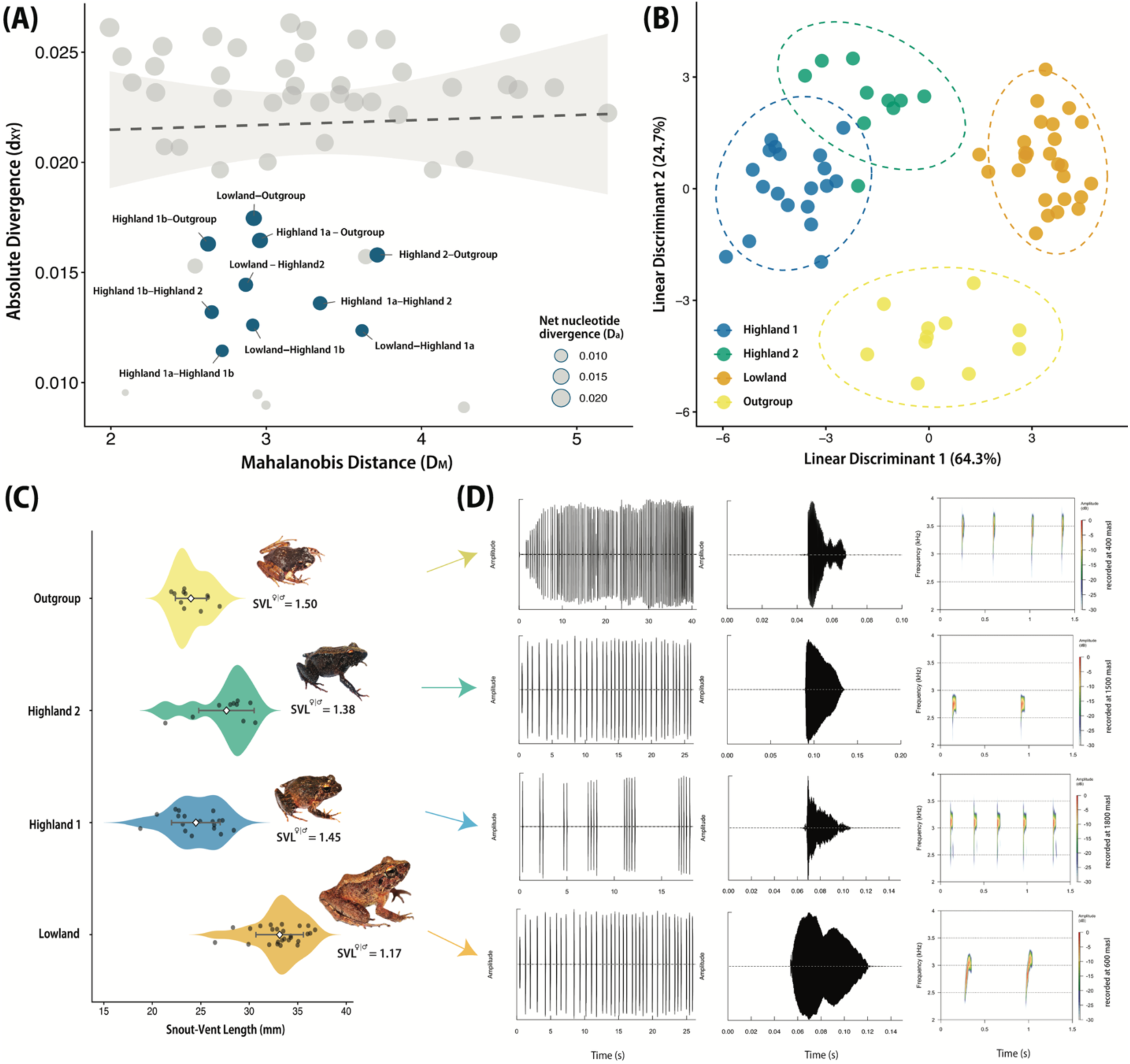
Morphological and behavioral divergence among focal lineages. **(A)** Absolute genomic divergence (*d_XY_*) plotted against Mahalanobis distance (*D_M_*), demonstrating that morphological differentiation does not scale predictably with genomic divergence among populations of Mindanao *Platymantis* (subgenus *Lahatnanguri*). **(B)** Linear Discriminant Analysis (LDA) illustrates clear multivariate separation among focal lineages, with minimal overlap between the two Highland lineages. **(C)** Violin plots of snout–vent length (SVL); sexual size dimorphism (female:male SVL ratio) is also shown. **(D)** Advertisement call structures of representative individuals; panels show, from left to right, a complete call oscillogram, an individual note oscillogram, and a representative 1.5-s spectrogram, illustrating divergence in temporal and spectral call properties among focal lineages.

The greatest morphological divergence (*D_M_*) was observed between the parapatric Highland 1a (BMR, northern slope) and the Lowland lineage (*D_M_* = 3.62), followed by the comparison between Highland 1a and Highland 2 (*D_M_* = 3.35). Highland 1a was consistently among the most morphologically distinct groups, whereas Highland 1b (BMR, southern slope) exhibited lower pairwise distances to the remaining lineages (*D_M_* = 2.65–2.91). Despite their geographic proximity within the BMR, Highland 1a and Highland 1b also showed moderate morphological differentiation (*D_M_* = 2.72), indicating phenotypic structuring within the high-elevation assemblage within the same mountain range. Distances involving Highland 2 and Lowland were intermediate (*D_M_* = 2.87), and Lowland–Highland 1b comparisons remained similarly differentiated (*D_M_* = 2.91).

Linear discriminant analysis (LDA) recovered clear separation among focal lineages (Figure 4B). The first axis (LD1; 64.3% variance explained) primarily separated the Lowland lineage from Highland 1 and Highland 2, with the Outgroup occupying an intermediate position. Linear Discriminant 1 was driven mainly by tibia length (coefficient = 34.83), thigh length (11.75), snout–vent length (SVL; 10.99), and tympanum diameter (11.92), opposed by forearm (–19.20) and hand length (–18.23). The second axis (LD2; 24.7%) further separated the Outgroup and was associated positively with hand length (25.39), tibia length (20.48), SVL (10.89), and internarial distance (15.37), and negatively with head width (–15.01) and upper arm length (–13.01).

Overall, variation in body size, limb proportions, head shape, and tympanum dimensions explained most morphological differentiation. Univariate comparisons revealed weaker and partially overlapping patterns. Lowland individuals were generally larger (33.1 ± 2.45 mm) than Highland 1 (25.0 ± 2.21 mm) and Highland 2 (28.4 ± 1.92 mm), while the Outgroup was smallest overall (24.0 ± 1.60 mm) (Figure 4C). Sexual size dimorphism was reduced in Lowland (SVL F/M ≈ 1.17) relative to Highland and Outgroup populations (SVL F/M ≈ 1.38–1.50).

Advertisement calls also differed among focal lineages in both temporal and spectral traits (Figure 4D). All lineages produced single-pulsed notes, but call structure varied along an inverse relationship between repetition rate and note duration. The Outgroup and Highland 1 exhibited the fastest calls (3.25 and 2.67 notes/sec, respectively) and shortest note durations (20.16 ± 5 ms and 30.16 ± 10.4 ms, respectively), whereas Highland 2 and the Lowland lineage produced slower calls (1.08 and 0.90 notes/sec, respectively) with nearly double the note duration (60.0 ± 5.1 ms and 60.9 ± 4.26 ms, respectively). These temporal differences were accompanied by distinct note envelopes (Figure 4D). Highland 1 produces short notes with abrupt onsets that rapidly decay to silence, whereas the Outgroup exhibits a distinct multi-phasic envelope with a trailing resonance tail. In contrast, Highland 2 produces fuller, asymmetrical triangular-shaped notes that taper gradually toward silence, whereas the Lowland lineage is unique in exhibiting bimodal (double-peaked) notes separated by a pronounced central amplitude depression.

Spectral properties further differentiated lineages despite partial overlap in frequency ranges (spectrograms in Figure 4D). The Outgroup exhibited the highest dominant frequency (2.876 ± 0.696 kHz), whereas Highland 2 showed the lowest (2.196 ± 0.382 kHz). Highland 1 (2.508 ± 0.433 kHz) and Lowland (2.344 ± 0.824 kHz) occupied intermediate spectral space, although Highland 1 maintained slightly higher mean frequencies than Lowland. Notably, calls of the Lowland lineage are distinctly frequency-modulated, exhibiting an upward frequency sweep (i.e., frequency changes throughout the note) that was absent in Highland 1 and Highland 2 lineages. Together, these results indicate acoustic divergence among lineages and most pronounced with parapatric Lowland and Highland 1 lineages, concordant with observed morphological divergence, despite shallow genome-wide differentiation.

## DISCUSSION

### Hierarchical diversification across montane systems

Phylogenomic analyses of ground-dwelling *Platymantis* (*Lahatananguri*) frogs from the Mindanao biogeographic region revealed a hierarchical pattern of diversification operating across nested spatial scales. The Lowland lineage is phylogenetically nested within highland populations and forms a sister relationship with Highland 1, rendering the high-elevation assemblage paraphyletic. This topology indicates a history of divergence structured across scales, with deeper separation between the Busa Mountain Range (BMR) and the Melibengoy Mountain Complex (MMC), and more recent diversification within the BMR. Within this framework, Highland 1 represents the closest extant high-elevation relative of the Lowland lineage.

Patterns of genomic differentiation further support this interpretation. The Lowland–Highland 1 pair occupies the shallow end of the genomic divergence continuum and exhibits patterns typical of genomic divergence in parapatric systems, where divergence is concentrated in only a few genomic regions (Feder et al. 2013; Martin et al. 2013; Seehausen et al. 2017). Notably, northern-slope individuals of Highland 1 carry mitochondrial genes associated with the Lowland lineage despite clustering with Highland 1 in ddRAD phylogenomic tree. Because we found no evidence of contemporary hybridization among our samples, this apparent mito-nuclear discordance appears more consistent with incomplete lineage sorting than ongoing gene flow (DeRaad et al. 2023)—especially given that divergence between Highland 1 and Lowland lineages appears relatively recent. In contrast, allopatrically distributed lineages showed greater genomic differentiation, with divergence being stronger between populations occupying contrasting habitats (Lowland vs. Highland 2) and weaker between those occupying similar habitats (Highland 1 vs. Highland 2). Together, these patterns suggest that ecological divergence may amplify the effects of geographic isolation, accelerating genome-wide differentiation as documented in lake–stream cichlids (Weber et al. 2021) and threespine sticklebacks (Weber et al. 2017).

Demographic and molecular rate patterns further support independent evolutionary trajectories among focal lineages. The Lowland lineage exhibits significantly elevated substitution rates relative to both highland lineages, which evolve at comparable and slower rates. This rate asymmetry appears consistent with colonization of warmer, low-elevation environments, where higher temperatures have been hypothesized to accelerate molecular evolution by increasing metabolic activity and mutation accumulation (Wright et al. 2010; Puurtinen et al. 2016). The Lowland lineage also exhibits higher genomic diversity and predominantly negative Tajima’s *D* values, patterns that suggest population expansion following its hypothesized transition or expansion into lower elevations. Highland 1 similarly shows elevated genomic diversity and negative Tajima’s *D* despite being restricted to high elevations (1500–2000 m a.s.l.) within the BMR. Although this pattern is compatible with demographic expansion, it may alternatively reflect targeted positive selection for specific genes associated with the physiological and ecological adaptation of amphibians living in high-elevation environments (Sun et al. 2018; Christodoulides et al. 2024). Distinguishing between these alternatives will require locus-specific analyses beyond the genome-wide summary statistics presented here. In contrast, the lower genomic diversity, short external branches in phylogenetic tree, and predominantly positive Tajima’s *D* values observed in Highland 2 point to a demographic contraction. This signature aligns with the population’s isolation in the high-elevation forests of MMC, where cataclysmic volcanic disturbances, as historically documented (Delfin Jr. et al. 1997), likely contributed to population declines.

Together, these results support a hierarchical model of diversification in which deeper allopatric divergence first separated populations among montane systems, followed by more recent lineage splitting within mountains along elevational gradients. One plausible mechanism is that repeated Pleistocene climate-induced sea-level oscillations alternately isolated and reconnected montane habitats across Mindanao (Brown et al. 2013; Justiniano et al. 2015). During interglacial periods of elevated sea levels, a once widespread high-elevation ancestor may have become fragmented among mountain systems, with remnant populations persisting in low-elevation forest refugia within the continental fragment of present-day Mindanao Island (i.e., Outgroup). This scenario is consistent with the shared distribution of the Outgroup with the highland lineages, all of which occur on one side of the SCDL and share mitochondrial haplotypes. The biogeographic affinity of lineages occurring west of the SCDL—a hypothesized paleo-island precursor to present-day Mindanao Island—has been recovered previously in phylogenomic studies (*Occidozyga laevis* frogs, Chan et al. 2022; *Dicaeum hypoleucum* birds, DeRaad et al. 2025), but our study is the first to explore these evolutionary relationships in depth using integrative datasets. Subsequent environmental shifts within mountains may then have created opportunities for ecological divergence across elevational gradients within the BMR. Diversification in this system therefore appears to have proceeded sequentially across regional and local spatial scales.

### Phenotypic and behavioral divergence outpace genome-wide differentiation

Although Lowland and Highland 1 represent one of the most recently diverged lineage pairs within the radiation of ground-dwelling *Platymantis* (*Lahatnanguri*) in the Mindanao biogeographic subregion, their shallow genome-wide differentiation contrasts sharply with pronounced divergence in morphology and acoustic mate-recognition signals. Similar patterns of rapid phenotypic divergence despite limited neutral genomic differentiation have been documented in populations occupying adjacent but ecologically contrasting habitats (Smith et al. 1997, 2005; Nosil 2012; Weber et al. 2017, 2021; Johannesson et al. 2024), reinforcing the role of ecological differences in promoting divergence in organismal traits early in the speciation process (Schluter 1996, 2001; Orr & Smith 1998; Nosil 2012; Shafer & Wolfe 2013). In our system, marked phenotypic differentiation has accumulated between the sister-pair Highland 1 and Lowland lineages despite weak genome-wide divergence, indicating that organismal divergence has progressed disproportionately relative to neutral genomic differentiation.

This pattern of marked phenotypic divergence despite weak genomic differentiation is further supported by the absence of a significant relationship between morphological and genome-wide genetic distance (*D_M_* – *d_XY_* correlations non-significant across datasets), with substantial body-shape differences persisting across a narrow range of genomic differentiation and over small geographic distances (also see Weber et al. 2021). Morphological divergence was not driven by single diagnostic characters but appears to reflect coordinated shifts in body size, sexual size dimorphism, limb proportions, and tympanum dimensions. Notably, the Lowland lineage exhibits a distinct overall phenotype relative to both highland lineages, whereas Highland 1 and Highland 2 retain comparatively conserved morphology despite deeper phylogenetic separation. This stark morphological divergence in parapatry, which is greatest between the continuously distributed Highland 1a and Lowland, is expected for populations occupying different environments, as has been shown in other systems (Smith et al. 1997, 2005; Hendry et al. 2009; Weber et al. 2017, 2021; Johannesson et al. 2024).

Behavioral divergence among *Platymantis* (*Lahatnanguri*) lineages appears equally pronounced. Because acoustic communication plays a central role in mate recognition in frogs (Wells 1977, Duellman & Trueb 1994, Wells & Schwartz 2007; Wilkins et al. 2013; Köhler et al. 2017), divergence in advertisement calls may contribute to assortative mating and reinforce reproductive isolation despite shallow genome-wide differentiation. Although temporal and spectral call properties, such as note rate and dominant frequency, are considered evolutionarily labile and can vary substantially among populations (Serrano et al. 2020; Rasolonjatovo et al. 2022), spectral properties such as patterns of frequency modulation are generally considered relatively evolutionarily conservative, or stable, and are often interpreted as lineage-specific acoustic traits (Duellman & Trueb 1994; Wells & Schwartz 2007; Köhler et al. 2017). In this context, the conspicuous upward frequency sweep uniquely observed in the Lowland lineage suggests that communication signals diverged likely alongside ecological transitions across elevation. This interpretation is further supported by observations from the northern slope of the BMR, where Highland 1a and Lowland occur in elevational contact yet maintain distinct call structures and show no evidence of contemporary hybridization, suggesting that acoustic divergence contributes to reproductive isolation in syntopy.

Our inference of the mismatch between neutral genomic divergence and phenotypic or behavioral differentiation is broadly consistent with ecological and genic models of speciation, in which divergent selection initially affects a limited subset of loci associated with adaptation and reproductive isolation while much of the genome remains weakly differentiated (Wu 2001; Nosil et al. 2009; 2016). Under this scenario, genome-wide divergence may lag behind ecologically or sexually relevant traits, potentially obscuring biologically meaningful lineage separation when assessed using neutral markers alone (Toews et al. 2016; Recuerda et al. 2025). More broadly, our results highlight the multidimensional nature of speciation, in which genomic, phenotypic, and behavioral traits can evolve at different rates and along partially independent trajectories (Bolnick et al. 2023; Johanneson et al. 2024). Empirical studies illustrate this decoupling across diverse systems. In African cichlid lake–stream population pairs, morphological diversification varies continuously along a gradient of reproductive isolation, whereas genomic differentiation shows a more pronounced break, with reproductive isolation increasing with genome-wide divergence (Weber et al. 2021). In capuchino seedeaters, by contrast, pronounced divergence in phenotypes involved in assortative mating and reproductive isolation can be associated with relatively few genomic regions, resulting in weak genome-wide differentiation (Recuerda et al. 2025). These examples, among others (discussed in Hendry et al. 2009; Nosil et al. 2009; Bolnick et al. 2023), demonstrate that the genomic, phenotypic, and behavioral traits and reproductive isolation components of divergence need not proceed in lockstep. Together, they emphasize the value of integrative datasets for characterizing lineage divergence and evaluating reproductive isolation across multiple traits (Cadena & Zapata 2021; Bolnick et al. 2023; Cerca & Stelkens 2026).

### Elevational gradients can promote divergence within montane systems

Mountain systems are widely recognized as major drivers of diversification (Janzen 1967; Cadena et al. 2011; Keller et al. 2013; Rahbek et al. 2019b), most commonly through allopatric isolation among disjunct high-elevation habitats that function as sky islands (Knowles & Richards 2005; Sanguila et al. 2011; Toussaint et al. 2013; Justiniano et al. 2015; Everson et al. 2020). However, mountains also encompass steep environmental gradients across remarkably short geographic distances, creating opportunities for lineage divergence within geographically continuous landscapes (Cadena et al. 2011). Under such conditions, populations occupying different elevations may experience contrasting selective environments while remaining geographically connected, allowing ecological divergence to accumulate despite limited spatial separation. At the same time, complex montane topography may generate localized habitat discontinuities that isolate populations over similarly small spatial scales (microallopatry; Habel et al. 2012; Scherz et al. 2023). Distinguishing between these alternative mechanisms is challenging because both can produce similar phylogenetic and genomic signatures during early divergence. However, ecological divergence along an elevational gradient should leave a broader signature of concordant geographic, phenotypic, and behavioral differentiation despite geographic continuity, providing a means to distinguish parapatric divergence from fine-scale spatial isolation.

This prediction is supported by our results, which are more consistent with divergence across an elevational, or more broadly ecological, gradient than with fine-scale spatial isolation alone. Along the northern slope of the BMR, Highland 1 and Lowland populations occur in geographic continuity across elevation, with limited spatial overlap and no obvious physical barrier separating them. Yet, despite shallow genome-wide differentiation, the two lineages maintain discrete genomic clusters and exhibit pronounced morphological and acoustic divergence. The contrast with the allopatrically distributed highland lineages provides an additional line of evidence for an ecological contribution to divergence. If fine-scale geographic isolation alone (in the absence of divergent ecological selection) were driving diversification, comparable phenotypic divergence might be expected among geographically separated populations (e.g., Smith et al. 2005; Hendry et al. 2009; Weber et al. 2021; James et al. 2025), including the two highland lineages. Instead, the strongest morphological and acoustic divergence is concentrated in the Lowland lineage, whereas the allopatrically distributed Highland 1 and Highland 2 lineages remain broadly similar in morphology and advertisement calls despite their relatively deeper genomic divergence. This asymmetry suggests that phenotypic divergence is more closely associated with the ecological transition between high and low elevations than with geographic separation per se. Although we cannot entirely exclude a contribution from mutation-order speciation, in which divergence can arise through independent evolutionary changes under similar selective conditions (James et al. 2025), the concordance between elevational distributions and phenotypic divergence supports a role for ecological differentiation in promoting lineage divergence within a geographically continuous mountain system.

The patterns recovered in our *Platymantis* system differ from many previously reported examples of putative elevational parapatric speciation. Based on theoretical expectations and empirical studies of parapatric divergence, we predicted that elevationally replaced lineages should occur along a continuous environmental gradient (Bush 1975) and, ideally, with a limited range size and reduced dispersal ability (Gavrilets et al. 2000), be sister lineages (Moritz et al. 2000; Emerson 2002; Losos & Glor 2003), show weak genomic differentiation despite geographic continuity and opportunities for gene flow (Feder et al. 2013; Martin et al. 2013; Seehausen et al. 2017), and exhibit concordant divergence in phenotypic or behavioral traits associated with ecological adaptation and reproductive isolation (Smith et al. 1997; Orr & Smith 1998; Schluter 2001; Nosil 2012; Funk et al. 2016). Previous studies have tested these predictions in putative cases of elevational parapatric divergence, but evidence has often remained equivocal. In several vertebrate systems, elevationally replaced taxa are not sister lineages, weakening the inference of divergence within the same elevational system (Caro et al. 2013; Rakotoarison et al. 2019; Linck et al. 2020). Other systems show marked phenotypic divergence without sufficient evidence of restricted gene flow (McCormack & Smith 2008; Parker et al. 2020; Hinckley et al. 2022; Páez-Vacas et al. 2022) or insufficient evidence that divergence has progressed toward reproductive isolation (Funk et al. 2016; Vences et al. 2023). Our system satisfies these predictions through the integration of island-wide genomic sampling with phylogenomic, morphological, acoustic, and geographic evidence within a single framework. Collectively, these independent lines of evidence support divergence between closely related lineages along an elevational gradient within the BMR and provide a stronger empirical case for elevation-mediated parapatric speciation.

More broadly, our findings support a longstanding view that diversification in oceanic islands is driven not only by isolation among islands, but also by evolutionary splitting within islands themselves (Whittaker et al. 2008; Heaney et al. 2013). Early biogeographic hypotheses proposed that the geological complexity and environmental heterogeneity of the Philippine archipelago promote diversification across multiple spatial scales, particularly within paleoendemic radiations like in genus *Platymantis* (Brown & Alcala 1970; Diamond & Gilpin 1983; Heaney 1986). However, empirical support for this hypothesis has historically been inferred largely from broad phylogeographic patterns (Brown et al. 2013). Only recently, through extensive, elevationally representative genetic sampling (e.g., Meneses et al. 2026) and the integration of multiple lines of evidence, have we begun to uncover the previously underappreciated role of elevational divergence operating within individual mountain systems. Together, these findings suggest that the extraordinary biodiversity of the Philippine archipelago may result not only from repeated colonization and allopatric divergence among islands—and among mountains within islands—but also from repeated diversification within mountains themselves. These empirical results provide support for the biological plausibility of parapatric speciation within single mountain systems and highlight an underrecognized mechanism contributing to biodiversity in topographically complex oceanic archipelagos.

## DATA AVAILABILITY

Raw sequencing reads have been deposited in the NCBI Sequence Read Archive under BioProject accession number PRJNA1533437.

## ETHICS APPROVAL STATEMENT

Specimens were collected and euthanized following protocols approved by the University of Kansas Institutional Animal Care and Use Committee (Animal Use Statement No. 279-01 to KMEP and AUS 158-05 to RMB). Fieldwork was conducted with prior informed consent from the Protected Area Management Board of the Allah Valley Protected Landscape (PAMB-AVPL Resolution No. 20 Series of 2023) and the relevant municipal local government units with jurisdiction over the study areas (Maitum, Kiamba, Lake Sebu, and Tboli), and under Wildlife Gratuitous Permit No. 337 issued by the Philippine Department of Environment and Natural Resources Biodiversity Management Bureau.

## AUTHOR CONTRIBUTIONS

**KMEP**: Conceptualization, Data curation, Formal analysis, Funding acquisition, Investigation, Methodology, Project administration, Resources, Visualization, Writing - Original Draft, Writing - Review & Editing; **FHT**: Conceptualization, Data curation, Investigation, Writing - Review & Editing; **AJLS**: Conceptualization, Data curation, Investigation, Resources, Writing - Review & Editing; **AM**: Conceptualization, Funding acquisition, Project administration, Resources, Supervision, Writing - Review & Editing; **RMB**: Conceptualization, Project administration, Resources, Supervision, Writing - Review & Editing.

## ACKNOWLEDGEMENTS

Primary fieldwork was supported by The Rufford Foundation through grants awarded to K. M. E. Pitogo (Project Nos. 26752-1 and 42230-2), with additional support from the University of Kansas Biodiversity Institute and Idea Wild. Sequencing was supported by the BioGEM Program, funded by the National Science Foundation (RaMP Award No. 2319820), in support of F. H. Tesfazion. Additional specimens deposited at KU were collected under NSF DEB 0743491 and 1654388 to R. M. Brown. We thank Pola Geneva Bumanglag and Anson Tagtag of the DENR Biodiversity Management Bureau for facilitating our Memorandum of Agreement and Wildlife Gratuitous Permit to collect biological specimens, and DENR Region XII Regional Executive Director Felix Alicer for issuing the Wildlife Export Permit. We are also grateful to the Protected Area Management Board of the Allah Valley Protected Landscape for granting clearance, DENR-PENRO South Cotabato for providing logistical support during fieldwork, and to the Municipal Local Government Units of Lake Sebu, Tboli, Maitum, and Kiamba for granting access to field sites. Finally, we sincerely thank the Indigenous Peoples and local communities of our field sites for their hospitality and support, as well as Angelica Emit, John Faus Yata, Mc Gabriel Fungan, John Michael Zante, and Rufilyn Dua for their invaluable assistance during fieldwork.

